# A comparison between time-constrained counts and line transects as methods to estimate butterfly diversity and monitor populations in tropical habitats

**DOI:** 10.1101/2021.09.04.458959

**Authors:** Attiwilli Suman, Nitin Ravikanthachari, Krushnamegh Kunte

**Affiliations:** National Centre for Biological Sciences, Tata Institute of Fundamental Research, GKVK Campus, Bellary Road, Bengaluru 560065, India

**Keywords:** butterfly monitoring, butterfly conservation, population sampling methods, biodiversity inventories, citizen science

## Abstract

Long-term species monitoring programmes have revealed catastrophic insect population declines and disruption of biological communities that are contributing to biodiversity loss. Such discoveries have been possible because of standardised methods, such as line transects, of counting butterflies and other insects. However, line transects are not feasible in many tropical and mountainous habitats, so alternative methods must be explored. To tackle this issue, we devised time-constrained (30-min) counts and compared butterfly diversity as estimated through this method with that estimated through line transects in three tropical habitats in India (evergreen forest, dry deciduous forest and an urban woodland). We tested the efficacy of the two methods to sample species richness and abundance, as well as numbers of rare, endemic and specialist butterflies. We observed greater species richness, and more species of habitat specialists and endemics per sample in time-constrained counts in evergreen forest, but not in the other two habitats. Thus, time-constrained counts were more efficient at detecting species in the species-rich evergreen habitat. Apart from this difference, the two sampling methods captured similar levels of species richness and other measures of diversity. Our study thus shows that time-constrained counts is a suitable if not a superior alternative to line transects to conduct butterfly diversity surveys and population monitoring in complex tropical landscapes. Due to methodological flexibility and simplicity, this method may be particularly useful to study the impacts of climate change, habitat fragmentation and land use practices on butterfly conservation in populous and tech-ready tropical countries using citizen science frameworks.

## INTRODUCTION

Survey and monitoring of insects is fundamental to their conservation (Kim 1993; Clark and Samways 1996; Brown 1997; Rohr et al. 2007; Leather et al. 2008). Spatial distribution maps obtained through ecological surveys can inform conservation planning and layout of reserves by identifying biogeographic units and areas of high species diversity, endemism or rarity (Kremen et al. 1993). The baseline data can also be used to set up monitoring programmes and assess the impact of anthropogenic disturbance on insect populations. This is especially relevant in tropical areas, where diversity of insects is extremely high but resources are scarce to undertake detailed studies of individual species (Braschler at al. 2010, Lewis and Basset 2007). Butterflies are a good target taxon for recording and monitoring because their taxonomy is relatively well-known, and they are conspicuous and relatively easy to identify in the field as compared to other insects (Thomas 2005; Basset et al. 2013).

Long-term butterfly monitoring programmes have been remarkably successful in the United Kingdom and Europe (Thomas 2005; van Swaay et al. 2008; Schmeller et al. 2009; Brereton et al. 2011). Their objective is to assess regional and national-level trends in abundance of butterfly species (van Swaay et al. 2008). All such programmes employ spatially explicit transects—popularly known in butterfly monitoring schemes as ‘Pollard walks’—in which recorders walk the transects at a uniform pace, counting all butterfly individuals observed within a certain width along the transect line (Pollard and Yates 1994). The transects are predefined, i.e., fixed a priori, and revisited every sampling session. The transect method is popular because it is simple to implement in open, relatively flat habitats with fewer butterfly species and lower population densities, and are easily replicable once the line transects have been identified by experienced butterfly watchers. The protocol allows rapid collection of data on relative abundances of butterflies using a fixed effort. Long-term population monitoring using this method has yielded valuable insights into population and community dynamics of British and European butterflies, including changes in community structure and diversity, and the recent alarming population declines, due to climate change, habitat fragmentation, and habitat alterations from land use practices (Roy and Sparks 2000; Van Dyck et al. 2009; Devictor et al. 2012; Botham et al. 2015; Thackeray et al. 2016; McDermott Long et al. 2017; Mills et al. 2017; van Strien et al. 2019; Warren et al. 2021a; Fourcade et al. 2021). Monitoring schemes of comparable magnitude are lacking in tropical countries and habitats, even though the tropics harbour a much larger proportion of butterfly diversity (Bonebrake et al. 2010).

The difficulty is that line transects and related statistical methods for data analysis were originally devised to count large mammals and other conspicuous organisms in simpler habitats. These methods have inherent limitations in monitoring insects in more complex habitats (Spitzer et al. 1993; Zhang et al. 2018). For example, line transects require the transect line to be more or less straight, which is nearly impossible to achieve in hilly tropical regions with dense vegetation, sometimes with difficult terrain. Moreover, line transects require recorders to cover the transect length at a uniform pace. Since butterflies and certain other insects are often highly clustered, e.g., at puddling sites in hundreds or thousands, the requirement of uniform pace is easily violated. Such highly clustered occurrence is also not easy to account for in the analysis of line transect data. Besides, the fixed-transect method has been criticized for poor detectability of cryptic, rare or sedentary species (Kadlec et al. 2012; Henry et al. 2015). Certain tropical butterfly genera are difficult to identify on the wing or may be easily missed while walking transects, especially in shaded areas (Walpole and Sheldon 1999; Caldas and Robbins 2003). However, line transects have been conducted at certain sites in tropical rainforests as part of previously established permanent vegetation monitoring plots in slightly more comfortable terrain (Basset et al. 2013).

There is a pressing need to develop appropriate protocols for sampling and monitoring of tropical butterflies. The pace of habitat destruction in the tropics is alarming, and at the same time interest in butterfly species discovery, diversity documentation, and conservation is on the rise. Individual researchers and institutions in the tropics may have adequate financial resources or manpower to conduct biodiversity assessments at small to medium spatio-temporal scales. However, citizen science—in conjunction with mobile apps, networking technologies and web platforms—is emerging as an important new movement in large-scale data generation for biodiversity research. Well-coordinated citizen science programmes using appropriate methods can provide large datasets of high quality that can contribute to conservation research and planning. Line transects, with their design requirements and reliance on costly preparations may not be amenable to be taken up at large scales in complex landscapes and/or by large groups. Alternative methods need to: (a) combine rigour with efficiency, (b) be relatively simple and cost-effective, and (c) be flexible enough to be implementable in a variety of terrains and habitat-types so that the protocols may be scaled up to regional and national levels. Initiatives such as day- or week-long national butterfly surveys, butterfly races, etc., have found alternatives that engage the general public, but they often lack uniform effort and rigour that are critical for the data to be comparable in a scientific analysis.

Time-constrained counts is an alternative method to line transects for conducting butterfly surveys and population monitoring (Beneš et al. 2003; Kadlec et al. 2012; Taron and Ries 2015; Kral-O’Brien et al. 2021). In time-constrained counts, effort is standardised by time rather than by length of the transect and area under the belt transect. It involves walking a wandering, often a zig-zag path rather than a fixed (more or less) straight line, thoroughly searching an area in the stipulated time, with the goal of maximizing the number of species and individuals observed (Fig. 1). This is accomplished by visiting resource patches—puddling sites, canopy gaps, roosting sites, etc., that are frequented by a diversity of butterfly species—and recording every individual of every species that is encountered. Unlike line transects, the survey path need not be fixed a priori, so it offers more flexibility for rapid surveys at new or even well-established field sites. Time-constrained counts also offer flexibility to navigate difficult terrain because a uniform pace and linear paths are not required, but they can still record spatial and temporal distribution of butterflies or their resources. Time-constrained counts may be especially advantageous in complex, dynamic landscapes, and even in smaller habitat fragments (Taron and Ries 2015). Similar to line transects, time-constrained counts assume that: (a) detectability is certain on the path, (b) counts are exact, (c) variation in the number of individuals counted at different sites or in different seasons truly reflect differences in their population sizes, and (d) butterflies are detected at their initial location (Taron and Ries 2015; Glennie et al. 2015). Other assumptions and violations of line transects should also apply, e.g., (a) individual organisms are not going back and forth on sampling paths, therefore they are likely to be counted only once, and (b) species with certain biological characteristics such as crepuscular activity patterns and excellent camouflage, are under-represented. It may be possible to apply some of the statistical methods and models originally developed for data generated through line transects to those generated through time-constrained counts. However, this needs to be specifically tested, and perhaps new statistical models and methods need to be developed for the time-constrained count method as the method becomes more popular with both research scientists and citizen scientists, and large-scale data become available.

**Fig. 1:**
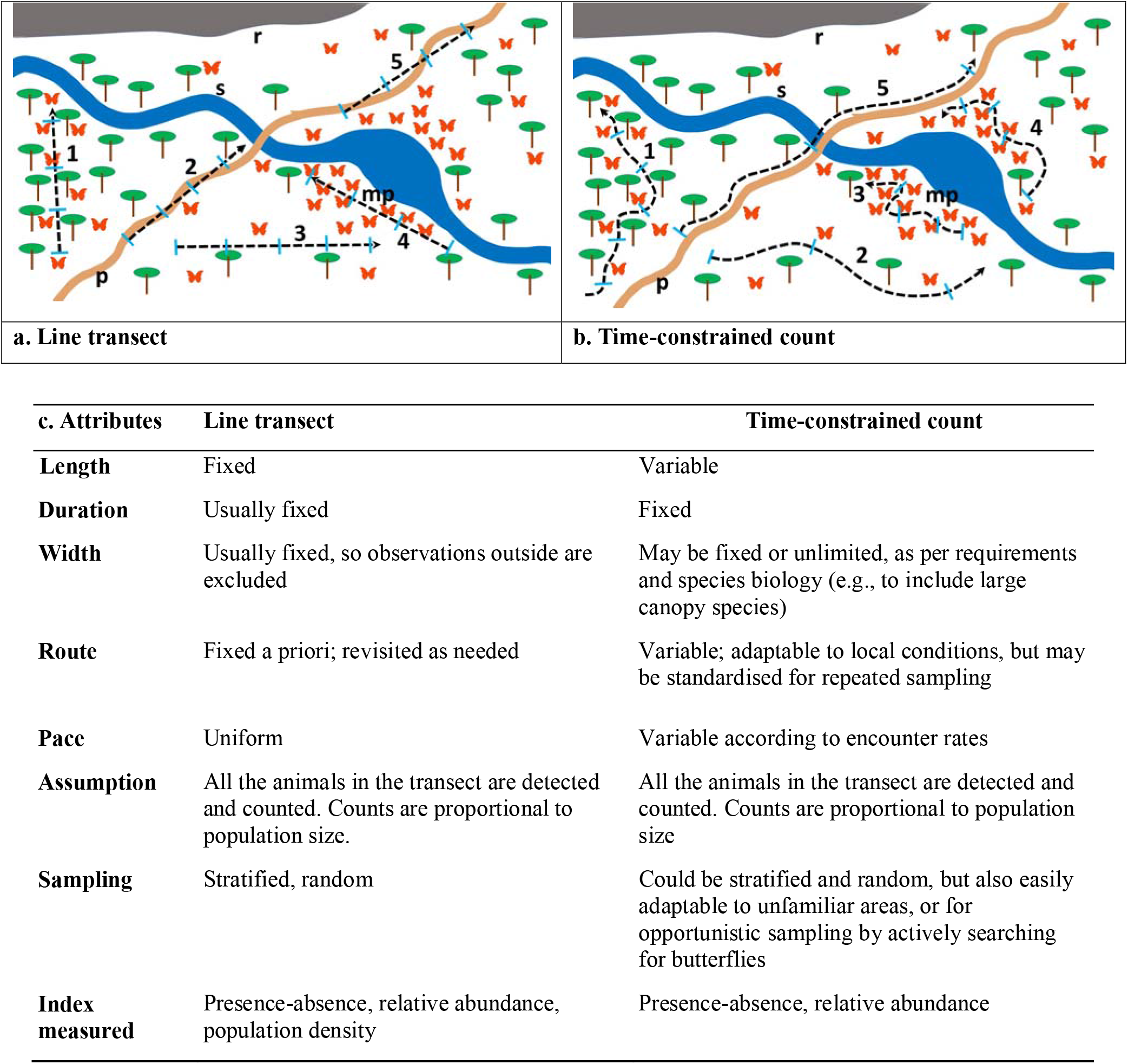
Schematic of a landscape, visualizing the two methods for conducting butterfly surveys i.e., line transect (a), and time-constrained count (b). Transects are usually straight lines laid near paths (*p*) and other landscape features, requiring walking at a uniform pace, during which organisms are sampled in a fixed area during a fixed timespan. The objective of time-constrained counts is to actively search an area for butterflies 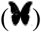, often sampling different micro-habitats, such as vegetation, rocky outcrops (r), stream-sides (s) and mud-puddling spots (mp), while keeping a fixed length of time as a unit of sampling but not the area. In a time-constrained count, a surveyor would spend more time in a patch where butterfly density is higher, whereas in a line transect the surveyor would normally be expected to maintain pace irrespective of butterfly density. As a result, the transect length remains more or less the same irrespective of butterfly density, whereas the physical length of a time-constrained count may be negatively correlated with butterfly density. A comparison of the attributes of line transects and time-constrained counts is given below (c).

Our study was motivated by a need to develop and test standardised methods for conducting butterfly surveys in tropical areas such as India, with its complex landscapes and high butterfly diversity. While India is home to four globally recognized biodiversity hotspots and over 1,400 butterfly species, large-scale data on the population dynamics of butterflies through seasons and space are scarce, even as anthropogenic pressures on butterfly habitats are mounting. This situation is widespread in other tropical countries as well. At the same time, with the rise in popularity of social media and citizen science initiatives, there is renewed interest in natural history, and an opportunity to engage enthusiastic naturalists to obtain large-scale data on butterfly diversity using standardised protocols. Towards this end, we compared two methods for butterfly surveys (line transects and time-constrained counts) in three habitats: (1) an evergreen forest, (2) a dry deciduous forest, and (3) an urban woodland, in southern India. We evaluated the efficacy of these methods to record the number of species and individuals as well as numbers of rare, endemic or specialist species.

## MATERIALS AND METHODS

### Survey methods

We conducted butterfly surveys in three habitats (evergreen forest, dry deciduous forest, urban woodland) (Fig. S1). All three study sites were located in Karnataka state of southern India (Fig. S1). The evergreen forest site (Brahmagiri Wildlife Sanctuary, 181.8 km^2^, altitude 100-1,600 m) is hilly with evergreen and semi-evergreen forests at low and mid-elevations, and grasslands and montane forest patches at higher altitudes. It is a protected area located in the Western Ghats biodiversity hotspot. The dry deciduous forest site (Savandurga State Reserve Forest, 26.6 km^2^, altitude 750-900 m) is located on the Dakhan (=Deccan) plateau. The urban woodland (Doresanipalya Forest Campus, 0.4 km^2^, altitude ~ 900 m), a reserved forest in the midst of the metropolitan city of Bengaluru, has a mix of native and exotic trees and mixed understorey, criss-crossed by paths used by citizens for morning walks.

The three sampling locations have been extensively surveyed for butterfly diversity over the past 11 years, with their butterfly faunas enumerated by a collaborative of professional and citizen scientists (Kunte and Ravikanthachari 2020; Kunte et al. 2020). Building on this robust experience, we wanted to compare the efficacy of the two survey methods in recording species and individuals in representative habitat patches within the three sites. Accordingly, we demarcated survey areas near 19 km and 10 km stretches of roads and forest paths at the evergreen and dry deciduous forest sites, respectively. The urban woodland was small and fully accessible and most of it was covered with both the survey methods. At each of the study sites, we surveyed butterflies using two methods for comparison: a) line transects (500 m in length, more or less in a straight line, 10 m in width, 10 m in height, covered in 10 mins), and b) time-constrained counts (length not fixed, covered in 30 mins) (Taron and Ries 2015) (Fig. 1). A single surveyor (NR) with 10 years of experience of watching and sampling butterflies at these locations, walked the line transect and count trails at a steady pace, noting down all the individuals of each species observed. We used GPS (Garmin eTrex Vista HCX) to record the location of the line transects, which were laid near forest trails (Fig 1). In the 30-min time-constrained counts, the surveyor (NR) walked a wandering path depending on the accessibility of terrain, adjusting the pace to the encounter rates of butterflies or their resources, often lingering around host plants, forest streams or canopy gaps that are frequented by butterflies (Fig. 1). To equalize the effort between the two methods, we walked three 10-min line transects immediately before or after every 30-min time-constrained count. We alternated between the two methods in each sampling session and in each habitat patch so that habitat variability did not affect results of a single method.

We carried out surveys between 0900 and 1300 hrs on clear, sunny days. We visited the sampling locations for 2–3 days each month and repeated the sampling four times at each location from February to May 2018 (Table 1). The total sampling effort for line transects and time-constrained counts for each habitat is shown in Table 1. The species recorded, their habitat-wise occurrence and conservation values are given in supplementary Table S1. Sampling datasets will be made available on Dryad upon provisional acceptance of the paper.

**Table 1:** Total monthly sampling effort per habitat and method (line transect: 500 m, 10 min; time-constrained counts: 30 min). Evergreen and dry deciduous forests were sampled monthly from Feb-May, the urban site was sampled from March-May.

|  | **Time-constrained**<br>**counts as, total (paths-**<br>**puddling spots)** | **Line transects** |
| --- | --- | --- |
| **Evergreen forest** |  |  |
| Feb | 7 (3-4) | 21 |
| Mar | 9 (4-5) | 24 |
| Apr | 8 (4-4) | 24 |
| May | 8 (4-4) | 24 |
| **Dry deciduous forest** |  |  |
| Feb | 7 | 21 |
| Mar | 7 | 21 |
| Apr | 6 | 18 |
| May | 6 | 18 |
| **Urban woodland** |  |  |
| Mar | 6 | 18 |
| Apr | 6 | 18 |
| May | 6 | 18 |

### Statistical analysis

We used rarefaction to compare the overall number of species observed via time-constrained counts vs. line transects. We used iNext package (Hsieh et al. 2016) in R 3.4.2 (R core team 2017) to compute rarefaction curves with 95% confidence intervals. The sample size-based rarefaction curves with bootstrapped confidence intervals can be used to quantify and compare species richness observed between samples (Chao et al. 2014). We pooled the data from the three 10-min line transects walked immediately before or after the time-constrained counts, and used 30 min as a uniform sampling effort to compare results of the two sampling methods. We computed rarefaction curves for the combined data from the four months of the survey. We used generalized linear models (GLMs) to compare number of species and number of individuals of butterflies recorded per sample (i.e., per 30 mins of line transect or time-constrained count) by the two methods (METHOD: *time-constrained counts* vs. *line transect*) in each of the three habitats, with survey month as a co-explanatory variable. For further analysis, we classified butterfly species encountered at the three study sites on the basis of endemism (Western Ghats endemics), rarity (rare or uncommon in the Western Ghats) and habitat specialisation (habitat preference for the endangered evergreen, semi-evergreen, riparian moist deciduous forests, and montane forests and grasslands), and assigned conservation value to each species as calculated by Kunte (2008). We ran regression models on the number of habitat specialists, endemics and rare species per sample similar to above with METHOD and MONTH as explanatory variables. Since both response variables were counts (of either individuals or species) we used poisson regression models with log-link function and negative binomial in case of overdispersion (Table 3–4). Since the dataset of endemics and rare species had excess of zero observations, we ran zero-inflated poisson (ZIP) models using the pscl package in R. The zero-inflated poisson model has two parts: a poisson count model and a logit model for predicting excess zeros. It is thus appropriate to model count data that have an excess of zero counts. We used non-parametric, two-sample Wilcoxon signed-rank test to compare conservation values of species observed in pairs of sampling methods in each of the three habitats.

**Table 2:** Total number of species, unique species (observed only in that habitat or method), habitat specialists, endemics and rare species observed via line transects and time-constrained counts at the evergreen forest, dry deciduous forest and urban woodland sites. Measurements for puddling spots are also shown for the evergreen site. Number of individuals recorded are shown in parentheses after the species numbers.

|  | **Total no. of**<br>**species**<br>**(individuals)** | **Unique**<br>**species** | **Habitat**<br>**specialists** | **Endemics** | **Rare species** |
| --- | --- | --- | --- | --- | --- |
| **Evergreen forest** | 140 (2722) | 69 (881) | 71 (1297) | 11 (57) | 25 (110) |
| Line transects | 109 (1080) | 19 (23) | 49 (517) | 7 (22) | 15 (48) |
| Time-constrained counts | 121 (1642) | 38 (85) | 62 (780) | 11 (35) | 19 (62) |
| Paths | 99 (677) | 9 (11) | 35 (307) | 4 (11) | 6 (17) |
| Puddling spots | 112 (965) | 22 (45) | 55 (473) | 10 (24) | 17 (45) |
| **Dry deciduous forest** | 106 (2428) | 21 (101) | 16 (155) | 0 (0) | 4 (7) |
| Line transects | 87 (1198) | 13 (27) | 10 (72) | 0 (0) | 4 (5) |
| Time-constrained counts | 87 (1230) | 13 (34) | 12 (83) | 0 (0) | 2 (2) |
| **Urban woodland** | 69 (806) | 4 (5) | 10 (54) | 0 (0) | 0 (0) |
| Line transects | 51 (404) | 12 (18) | 4 (24) | 0 (0) | 0 (0) |
| Time-constrained counts | 57 (402) | 18 (19) | 10 (30) | 0 (0) | 0 (0) |

**Table 3:** Estimated regression parameters, standard errors, z-values and P-values for the GLMs of the abundance and number of species observed per sample by the two methods, viz. line transects and time-constrained counts at the three habitats, with month of survey (season) as a covariate. Intercept represents line transects in February for evergreen and dry deciduous forest and March for urban woodland.

| Species per sample |  |  |  |  | Abundance per sample |  |  |  |  |
| --- | --- | --- | --- | --- | --- | --- | --- | --- | --- |
|  | Variable | Estimate | Std. error | z value | P-value | Estimate | Std. error | z value | P-value |
| Evergreen forest |  | (negative binomial) |  |  |  | (negative binomial) |  |  |  |
|  | Intercept | -0.51 | 0.09 | -5.71 | < 0.01 | 0.87 | 0.15 | 5.92 | < 0.01 |
|  | Time-constrained counts | 0.08 | 0.08 | 0.94 | 0.35 | 0.24 | 0.12 | 1.91 | 0.06 |
|  | March | 0.05 | 0.11 | 0.45 | 0.65 | -0.24 | 0.19 | -1.26 | 0.21 |
|  | April | -0.47 | 0.11 | -4.13 | < 0.01 | -1.07 | 0.18 | -6.04 | < 0.01 |
|  | May | -0.56 | 0.11 | -4.94 | < 0.01 | -1.26 | 0.17 | -7.24 | < 0.01 |
| Dry deciduous forest |  | (poisson) |  |  |  | (negative binomial) |  |  |  |
|  | Intercept | 2.58 | 0.08 | 31.82 | < 0.01 | 3.4 | 0.1 | 32.51 | < 0.01 |
|  | Time-constrained counts | -0.04 | 0.07 | -0.58 | 0.57 | 0.02 | 0.09 | 0.21 | 0.84 |
|  | March | 0.26 | 0.11 | 2.46 | 0.01 | 0.42 | 0.14 | 3 | < 0.01 |
|  | April | 0.31 | 0.09 | 3.32 | < 0.01 | 0.65 | 0.12 | 5.22 | < 0.01 |
|  | May | 0.43 | 0.1 | 4.33 | < 0.01 | 0.54 | 0.13 | 4.01 | < 0.01 |
| Urban woodland |  | (negative binomial) |  |  |  | (negative binomial) |  |  |  |
|  | Intercept | 2.26 | 0.12 | 18.67 | < 0.01 | 3.16 | 0.17 | 18.64 | < 0.01 |
|  | Time-constrained counts | -0.02 | 0.12 | -0.18 | 0.86 | 0.01 | 0.17 | 0.06 | 0.96 |
|  | April | -0.06 | 0.15 | -0.42 | 0.68 | -0.32 | 0.21 | -1.54 | 0.13 |
|  | May | 0.23 | 0.14 | 1.58 | 0.11 | 0.12 | 0.21 | 0.57 | 0.57 |

**Table 4:** Estimated regression parameters, standard errors, z-values and P-values for the GLMs of the abundance and number of species of habitat specialists observed per sample by the two methods, viz. line transects and time-constrained counts in the three habitats, with month of survey (season) as a covariate. The same analysis was repeated for rare and endemic species at the evergreen forest site using zero-inflated poisson (ZIP) regression. Intercept represents line transects in February for evergreen and dry deciduous forests and March for urban woodland.

|  |  | Species per sample |  |  |  | Abundance per sample |  |  |  |
| --- | --- | --- | --- | --- | --- | --- | --- | --- | --- |
| Variable |  | Estimate | Std. error | z value | P-value | Estimate | Std. error | z value | P-value |
| *Habitat specialists* |  |  |  |  |  |  |  |  |  |
| **Evergreen forest** |  | (negative binomial) |  |  |  | (negative binomial) |  |  |  |
|  | Intercept | 2.02 | 0.14 | 14.85 | < 0.01 | 3.32 | 0.21 | 16.15 | < 0.01 |
|  | Time-constrained counts | 0.27 | 0.12 | 2.31 | 0.02 | 0.35 | 0.17 | 2.02 | 0.04 |
|  | March | -0.01 | 0.17 | -0.05 | 0.96 | -0.13 | 0.26 | -0.51 | 0.61 |
|  | April | -0.51 | 0.17 | -3.05 | < 0.01 | -0.95 | 0.25 | -3.86 | < 0.01 |
|  | May | -0.39 | 0.16 | -2.45 | 0.01 | -0.87 | 0.24 | -3.65 | < 0.01 |
| **Dry deciduous forest** |  | (poisson) |  |  |  | (negative binomial) |  |  |  |
|  | Intercept | -0.06 | 0.3 | -0.21 | 0.83 | -0.09 | 0.33 | -0.26 | 0.79 |
|  | Time-constrained counts | -0.02 | 0.22 | -0.11 | 0.91 | 0.16 | 0.23 | 0.7 | 0.49 |
|  | March | 0.66 | 0.36 | 1.82 | 0.07 | 1.14 | 0.39 | 2.92 | < 0.01 |
|  | April | 0.7 | 0.33 | 2.12 | 0.03 | 1.49 | 0.35 | 4.24 | < 0.01 |
|  | May | 0.77 | 0.34 | 2.23 | 0.03 | 1.18 | 0.38 | 3.15 | < 0.01 |
| **Urban woodland** |  | (poisson) |  |  |  | (negative binomial) |  |  |  |
|  | Intercept | -0.56 | 0.44 | -1.26 | 0.21 | -0.27 | 0.45 | -0.6 | 0.55 |
|  | Time-constrained counts | 0.22 | 0.37 | 0.6 | 0.55 | 0.14 | 0.39 | 0.36 | 0.72 |
|  | April | 0.12 | 0.49 | 0.24 | 0.81 | 0.09 | 0.51 | 0.18 | 0.86 |
|  | May | 0.81 | 0.42 | 1.91 | 0.06 | 1.19 | 0.47 | 2.66 | < 0.01 |
| *Rare species (evergreen site)* |  |  |  |  |  |  |  |  |  |
|  | Count model coefficients (poisson) |  |  |  |  | Count model coefficients (poisson) |  |  |  |
|  | Intercept | 0.83 | 0.29 | 2.82 | <0.01 | 1.35 | 0.21 | 6.43 | <0.01 |
|  | Time-constrained counts | -0.02 | 0.3 | -0.07 | 0.94 | -0.12 | 0.21 | -0.57 | 0.57 |
|  | March | -0.03 | 0.33 | -0.1 | 0.92 | 0.07 | 0.25 | 0.29 | 0.77 |
|  | April | -0.43 | 0.47 | -0.92 | 0.36 | -0.32 | 0.29 | -1.09 | 0.27 |
|  | May | -1.42 | 0.38 | -3.74 | <0.01 | -1.29 | 0.46 | -2.8 | <0.01 |
|  | ZIP model coefficients (binomial) |  |  |  |  | ZIP model coefficients (binomial) |  |  |  |
|  | Intercept | -1.18 | 1.15 | -1.02 | 0.3 | -0.9 | 0.8 | -1.13 | 0.26 |
|  | Time-constrained counts | -1.31 | 1.39 | -0.95 | 0.34 | -0.87 | 0.79 | -1.11 | 0.27 |
|  | March | 0.31 | 1.44 | 0.22 | 0.83 | 0.15 | 1.02 | 0.15 | 0.88 |
|  | April | 1.37 | 1.44 | 0.95 | 0.34 | 1.41 | 0.91 | 1.54 | 0.12 |
|  | May | -9.71 | 212.62 | -0.05 | 0.96 | 0.22 | 1.65 | 0.13 | 0.9 |

Table 4. (contd.)
| *Endemic species* (evergreen site) |  |  |  |  |  |  |  |  |
| --- | --- | --- | --- | --- | --- | --- | --- | --- |
| Count model coefficients (poisson) |  |  |  |  | Count model coefficients (poisson) |  |  |  |
| Intercept | -0.56 | 0.46 | -1.22 | 0.22 | 0.75 | 0.35 | 2.14 | 0.03 |
| Time-constrained counts | 0.83 | 0.41 | 2.02 | 0.04 | 0.27 | 0.36 | 0.75 | 0.46 |
| March | -0.62 | 0.7 | -0.89 | 0.37 | 0.07 | 0.48 | 0.15 | 0.88 |
| April | -0.31 | 0.59 | -0.52 | 0.6 | -0.58 | 0.5 | -1.15 | 0.25 |
| May | -0.14 | 0.55 | -0.25 | 0.8 | -0.54 | 0.46 | -1.19 | 0.23 |
| ZIP model coefficients (binomial) |  |  |  |  | ZIP model coefficients (binomial) |  |  |  |
| Intercept | -10.66 | 251.79 | -0.04 | 0.97 | 0.51 | 0.72 | 0.71 | 0.48 |
| Time-constrained counts | 11.09 | 251.79 | 0.04 | 0.97 | -0.35 | 0.7 | -0.5 | 0.62 |
| March | -5.58 | 92.62 | -0.06 | 0.95 | 0.23 | 0.9 | 0.26 | 0.8 |
| April | -1.38 | 1.92 | -0.72 | 0.47 | -0.43 | 0.95 | -0.45 | 0.65 |
| May | -2.47 | 2.99 | -0.83 | 0.41 | -1.02 | 1.01 | -1.01 | 0.31 |

## RESULTS

Number of individuals, species, habitat specialists, endemics and rare species observed during the survey are summarized in Table 2. Measures of species diversity, endemism and other parameters relevant for butterfly sampling across the two sampling methods are analysed and compared below.

### Species richness

Species richness was similar in time-constrained counts and line transects in two habitats: urban woodland (time-constrained counts = 57 species, lower confidence limit (LCL) = 49.4, upper confidence limit (UCL) = 64.6; line transects = 51 species, LCL = 46.2, UCL = 55.8), and dry deciduous forest (time-constrained counts = 87 species, LCL = 79.3, UCL = 94.7; line transects = 87 species, LCL = 79.4, UCL = 94.7) (Fig. 2a-b). Species richness was higher in time-constrained counts as compared to line transects at the evergreen forest site (time-constrained counts = 122 species, LCL = 111.3, UCL = 133.5; line transect = 102 species, LCL = 93.8, UCL = 110.2) (Fig. 2c). This could be partly due to higher number of species observed in the time-constrained counts at mud-puddling spots and small openings along forest paths as compared to time-constrained counts and line transects along shaded forest paths (time-constrained counts at mud-puddling spots and forest openings = 92 species, LCL = 83.1, UCL = 101.3; time-constrained counts along forest paths = 81 species, LCL = 71.6, UCL = 90.4; line transects = 73 species, LCL = 66.6, UCL = 80.3) (Fig. 2d).

**Fig. 2:**
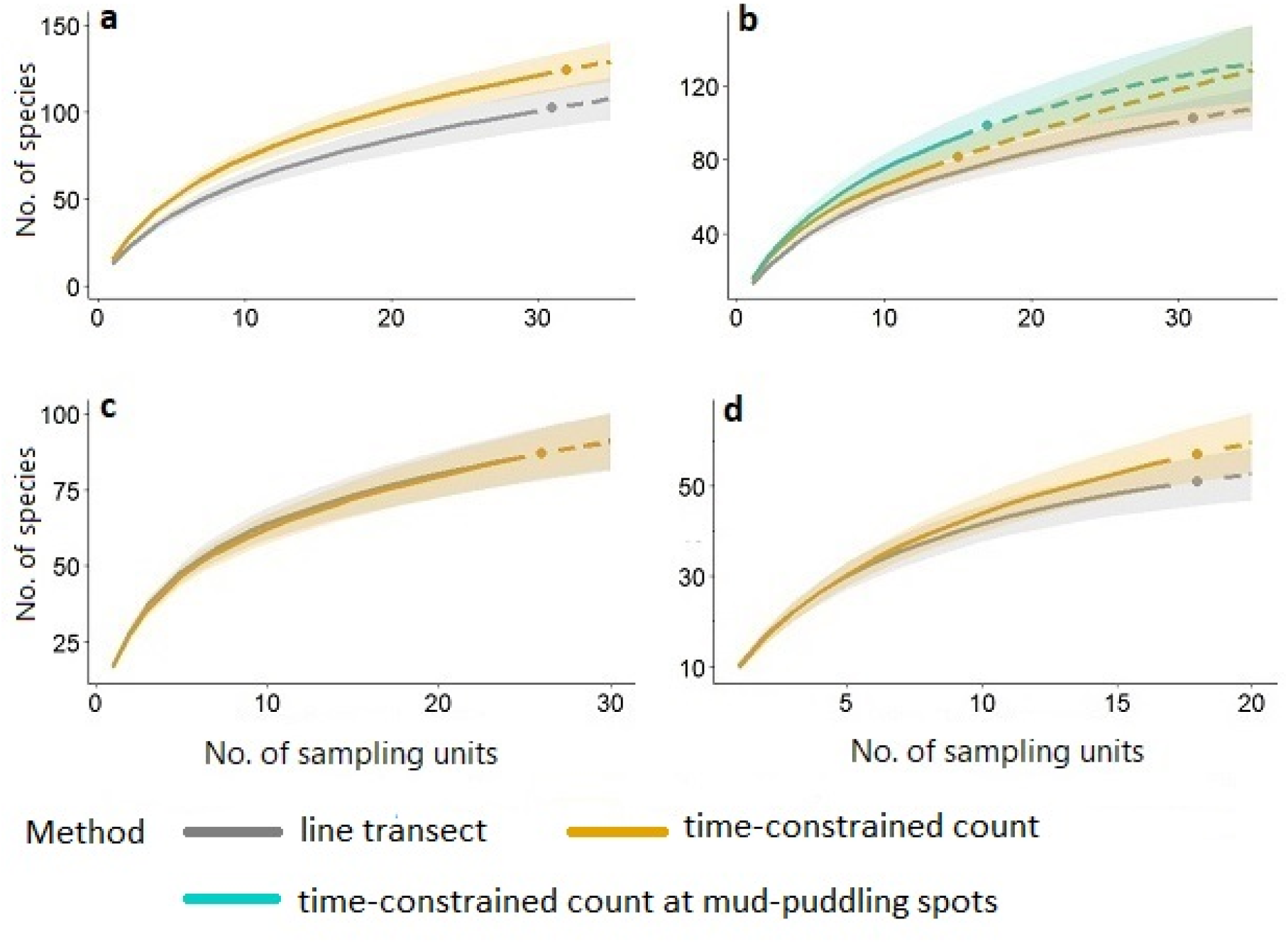
Rarefaction curves with 95% CIs, comparing the species richness observed in line transects and time-constrained counts un urban woodland (a), dry deciduous forest (b), and evergreen forest (c, d) habitats. Continuous lines = interpolation, stippled lines = extrapolation.

### Numbers of species and individuals per sample

Numbers of species and individuals per sample were similar between time-constrained counts and line transects in all three habitats (Table 3, Fig. 3). However, there was significant monthly variation in species and abundance per sample in the evergreen (species: *χ^2^*(3)=32.73, *p*<0.00; abundance: *χ^2^*(3)=49.71, *p*<0.00; Fig. 4; Table 3) and dry deciduous forests (species: *χ^2^*(3) = 20.57, *p*<0.00; abundance: *χ^2^*(3) = 22.76, *p*<0.00; Fig.4; Table 3). While the species per sample reduced from Feb–May in the evergreen forest, it increased during the same period in the dry deciduous forest habitat (Fig. 4; Table 3).

**Fig. 3:**
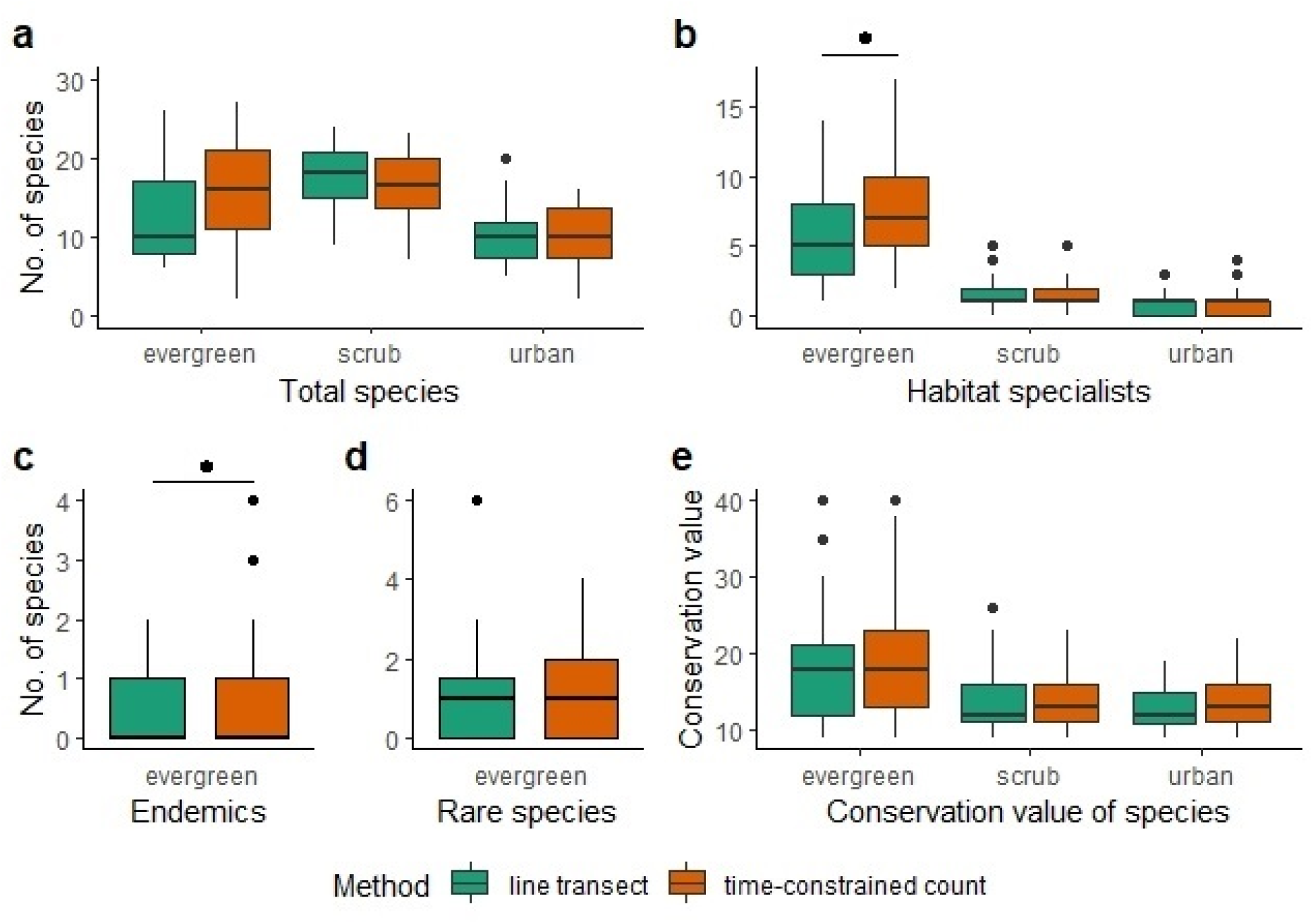
Boxplots showing total number of species (a), habitat specialists (b), endemics (c) and rare species (d) per sample, and conservation values of species (e) observed via two methods, viz. line transects and time-constrained counts in evergreen forest, dry deciduous forest and urban woodland.

**Fig. 4:**
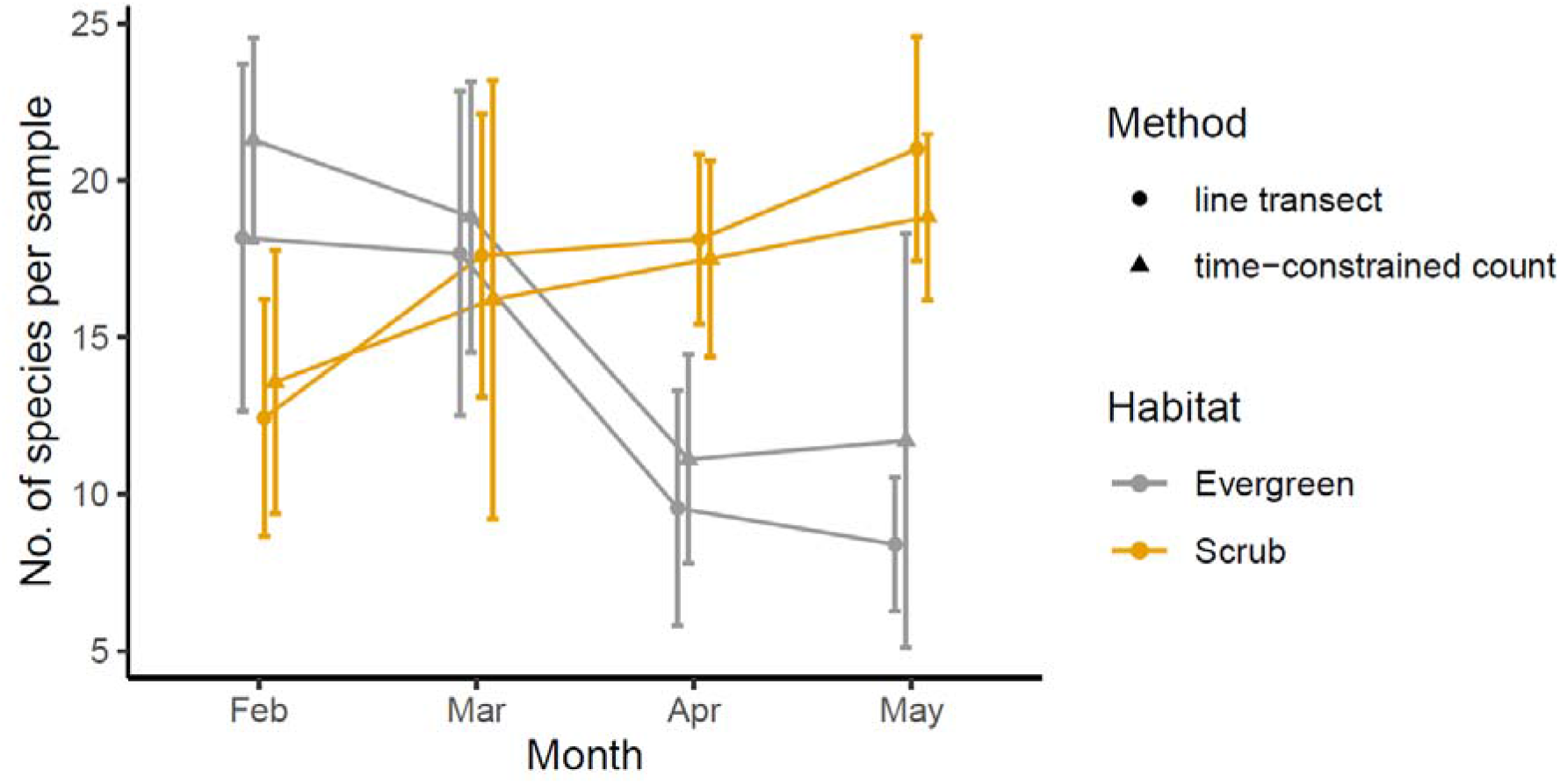
Monthly variation in number of species observed per sample in line transects and time-constrained counts in the evergreen and dry deciduous forests. While the species per sample reduced from Feb-May in the evergreen forest, it increased during the same period in the dry deciduous forest habitat.

### Habitat specialists, rare and endemic species

Numbers of species and individuals of habitat specialists observed *per sample* were higher in time-constrained counts only in the evergreen forest (no. of species: *Est*_timed_=0.27±0.12, *z*=2.31, *p*=0.02; no. of individuals: *Est*_timed_=0.35±0.17, *z*=2.02, *p*=0.04) (Table 4, Fig. 3). For rare and endemic species, since the overall abundance in the dry deciduous forest and urban woodland was extremely low (rare species: dry deciduous forest=7, urban woodland=0; endemic species: dry deciduous forest=0, urban woodland=0; Table 2), we conducted further analysis for these species only for the evergreen forest. A greater number of endemic species (but not their abundance) per sample was observed in time-constrained counts as compared to line transects (Poisson count model coefficients for no. of endemic species: *Est*_timed_=0.83±0.41, *z*=2.02, *p*=0.04), while no difference was found for rare species (Table 4, Fig. 3).

### Conservation value

We observed no difference in conservation values of species encountered by the two methods in any of the three habitats (Fig. 3). However, evergreen forest had a greater number and abundance of species with a high conservation value as compared to dry deciduous forest (Wilcoxon rank sum test: *W*=7880, *p*<0.00) and urban woodland (Wilcoxon rank sum test: *W*=5532.5, *p*<0.00), as revealed by both the survey methods (Fig. 3).

## DISCUSSION

Long-term butterfly monitoring programmes have generated valuable data on population dynamics and conservation status of butterflies (Warren et al. 2021b). Most of these studies have been in temperate countries in Europe. This paper was motivated by the need to develop inexpensive and easily implemented methods for monitoring butterflies in tropical and mountainous areas.

We compared two visual-count survey methods, namely, line transects and time-constrained counts, in three different habitats: (1) evergreen forest, (2) dry deciduous forest, and (3) urban woodland. We observed similar species richness via time-constrained counts and line transects in the urban woodland and dry deciduous forest. However, we observed greater species richness via time-constrained counts in the evergreen forest. We also observed greater number of species and abundance of habitat specialists, and greater number of species of habitat specialists and endemics *per sample*, in time-constrained counts as compared to line transects in evergreen forest. These results are parallel to those from other studies at different scales and habitats (Kadlec et al. 2012; Kral-O’Brien et al. 2021).

Species richness is a widely used index in ecology and conservation, but surprisingly difficult to measure accurately (Gotelli and Colwell 2001). Raw species counts underestimate species richness due to many species remaining undetected. Moreover, detectability varies between species and observers and over space, time and habitats (Kéry and Plattner 2007; Nowicki et al. 2008; Isaac et al. 2011; Pellet et al. 2012). Time-constrained counts may improve detectability of species by enabling the surveyor to search a habitat more thoroughly at a pace adapted for local habitat conditions or to species being sampled, and spending more time where butterflies may be concentrated. This may be especially true for rare, cryptic or specialist species, and those that are difficult to identify in flight. Our study detected higher number of habitat specialist and endemic species per sample in time-constrained counts in the more complex evergreen forest habitat. This was partly due to inclusion of puddling spots in the surveys, which sampled overall greater numbers of species, habitat specialists, endemics and rare species as compared to line transects (Table 2). Puddling spots attract concentrations of many species since they are a crucial resource for butterflies, including species that may be easily missed in line transects, such as smaller-sized, fast-flying or largely canopy species. On the other hand, the results of time-constrained counts vs. line transects were similar in less structurally complex habitats (dry deciduous forest and urban woodland), where the possibility of missing canopy species is smaller, and there are fewer species that can be identified only with close inspection. Thus, taking a non-linear path, paying particular attention to butterfly resources such as nectar/host plants, wet soil, tree sap, and rotting fruit, crabs or fish may indeed improve species detection and lead to better estimates of species richness in complex, species-rich habitats.

Traditionally, bait traps have been used commonly for butterfly surveys in tropical regions, e.g., to evaluate effects of natural disturbance such as tree-fall gaps (Hill et al. 2001; Pardonnet et al. 2013), vertical stratification of butterfly communities (Fermon et al. 2003; Molleman et al. 2006), forest fragmentation (Uehara-Prado et al. 2007), and intensification within agroecosystems (Mas and Dietsch 2003). However, bait-trapping is suitable to surveying a narrow range of subfamilies and genera within one family (Nymphalidae). This method is not suitable to sample larger butterfly communities, and this method can also not scale up over large areas. Hence, bait-trapping is complementary to visual census methods such as transect counts (Sparrow et al. 1994; Jakubikova and Kadlec 2015; Kral et al. 2018). Given that time-constrained counts are more efficient at detecting species and simpler to conduct in complex terrain, this method is particularly suited for studies requiring occupancy surveys and for mapping butterfly diversity at large scales. Since the survey path need not be fixed in time-constrained counts, recorders may vary in their search efficiency and attention paid to particular resources. Observer bias can be minimized by training surveyors prior to conducting the surveys, and by noting down GPS tracks of surveys. For ecological studies at smaller spatial scales, given better detectability via time-constrained counts as compared to line transects even at relatively small study plots (Kadlec et al. 2012), a grid-based, area-proportionate scheme of time-constrained counts by a team of well-trained recorders could potentially yield high-quality abundance-based data on butterflies. Such a method is also suitable for projects involving citizen scientists.

It should be noted that both line transects and time-constrained counts represent a measure of butterfly activity rather than true abundance. These methods also cannot be used to estimate population sizes without a lot more additional data generated through methods such as capture-mark-recapture. While line transects standardise effort by length of the transect, time-constrained counts do so by time expended. Thus, many of the methodological factors that influence counts of butterflies in line transects (including pollard walks, discussed in Basset et al. 2013) are equally applicable to time-constrained counts. For example, environmental factors such as ambient temperature, wind velocity and elevation, may need to be either controlled for during survey efforts (van Swaay et al. 2008), or measured and accounted for during statistical analysis (Basset et al. 2013). Seasonal variation can also considerably affect butterfly numbers. We found significant seasonal variation in butterfly abundance at our study sites. Thus, depending on the goals of the study, number of surveys per year will need to be optimized for adequate representation of seasonal variation (Basset et al. 2013). Similarly, for large areas, sampling locations must be representative of all the habitats in the area (Basset et al. 2013). For mapping and monitoring studies at local and/or large spatial scales, protocols for time-constrained counts can be formalised, so that the counts are replicable and comparable across space and time. These may include setting up tracks with GPS locations, time of day, sampling frequency, a predefined set of surveyors, repeated sampling, etc., formalised and agreed to by a research team for long-term monitoring. The strictness of the methodological criteria for monitoring should depend on the scope of the monitoring programme (Lindenmayer and Likens 2010), such as whether it is conducted at local or regional scales (Chiarucci et al. 2011; Scheiner et al. 2011), and whether professional or citizen scientists are involved in a formalised, centrally coordinated project. In general, more question-driven and goal-oriented monitoring programmes would require stricter design criteria. In any case, the time-constrained count method can be formalised and standardised just as well as line transects and other sampling methods for a given project, although it does appear to be more useful in tropical and mountainous terrains, as discussed above. For more question-driven and goal-oriented research projects, time-constrained counts may be used in conjunction with bait-trapping, capture-mark-recapture and other methods to estimate population sizes, community composition, and species diversity (Taron and Ries 2015).

Based on our findings as well as those of other recent studies (Kadlec et al. 2012; Kral-O’Brien et al. 2021), we conclude that time-constrained counts are a viable and perhaps even a superior alternative to line transects for conducting butterfly surveys and to monitor butterfly populations in tropical and mountainous habitats. Statistical methods to analyse data generated through time-constrained counts are not yet as well developed as those for transects, or they have limited statistical models available for estimation of population sizes and community diversity. However, many of the existing methods may be easily adapted. Considering the difficulty of conducting line transects in tropical and mountainous areas, and the need to develop alternative methods such as time-constrained counts, it will serve the scientific and conservation community to urgently develop these methods. This may prove to be especially important at a time when large amounts of data are being generated through citizen science projects. It is easier to train people in conducting time-constrained counts, and the counts are also easier to implement. So, such data may be more easily formalised for subsequent scientific analysis. Smartphones and citizen science platforms can greatly facilitate these steps.

## Acknowledgements

We thank Tarun Karmakar, Minoti Asher, Swathi H. A., Santosh Hatti, and Nagesh Gowda for assistance in the field, and Karnataka Forest Department for research permission (letter no. PCCF/WL/E2/CR/227/2014-2015 dated 26/05/2017) to carry out this study, for which we thank the Principal Chief Conservator of Forest (Wildlife) and field officers of the state. This work was funded by an NCBS Bridging Fellowship to SA, and a research grant from NCBS to KK.

## SUPPLEMENTARY MATERIAL

**Table S1.**
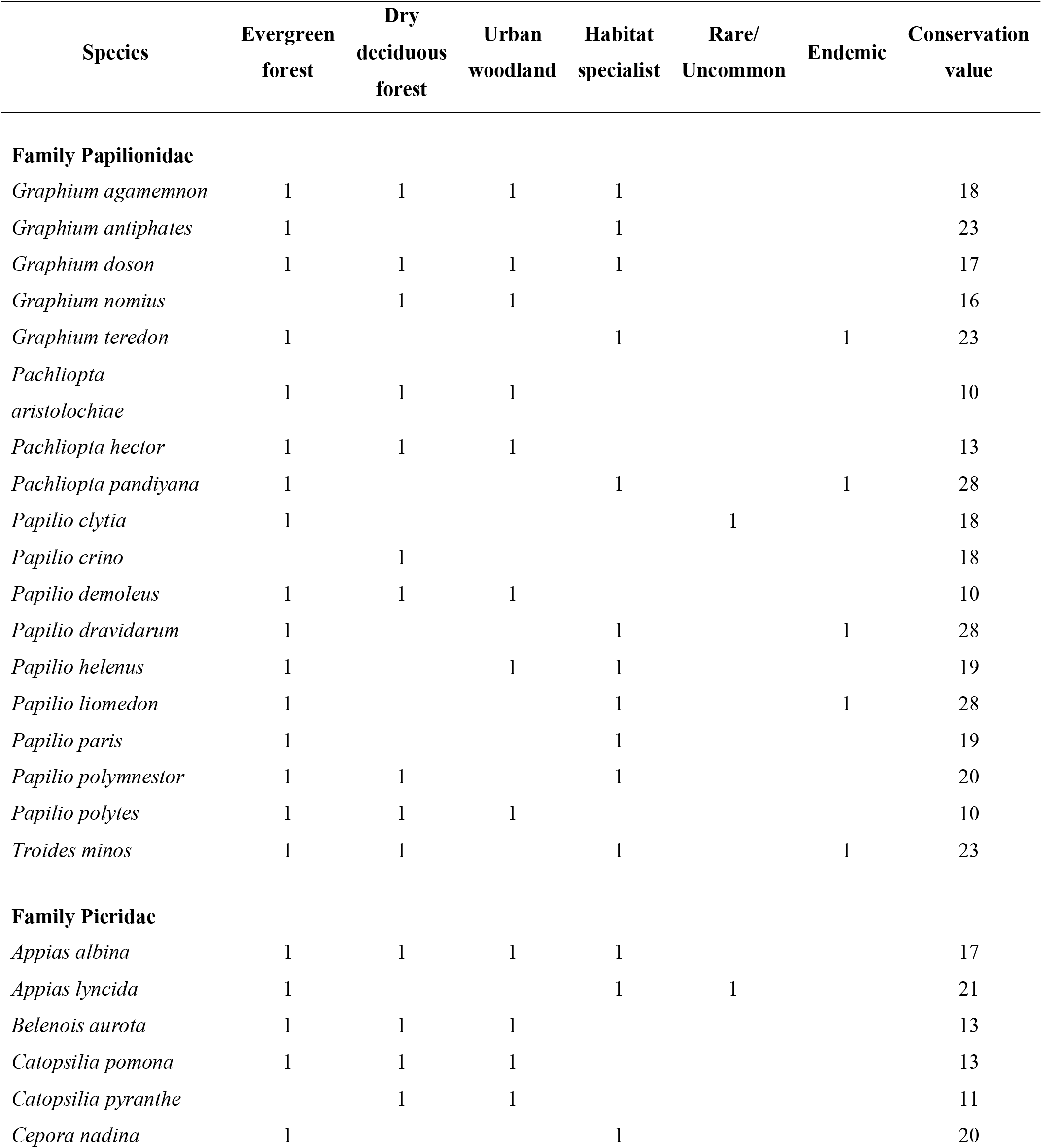
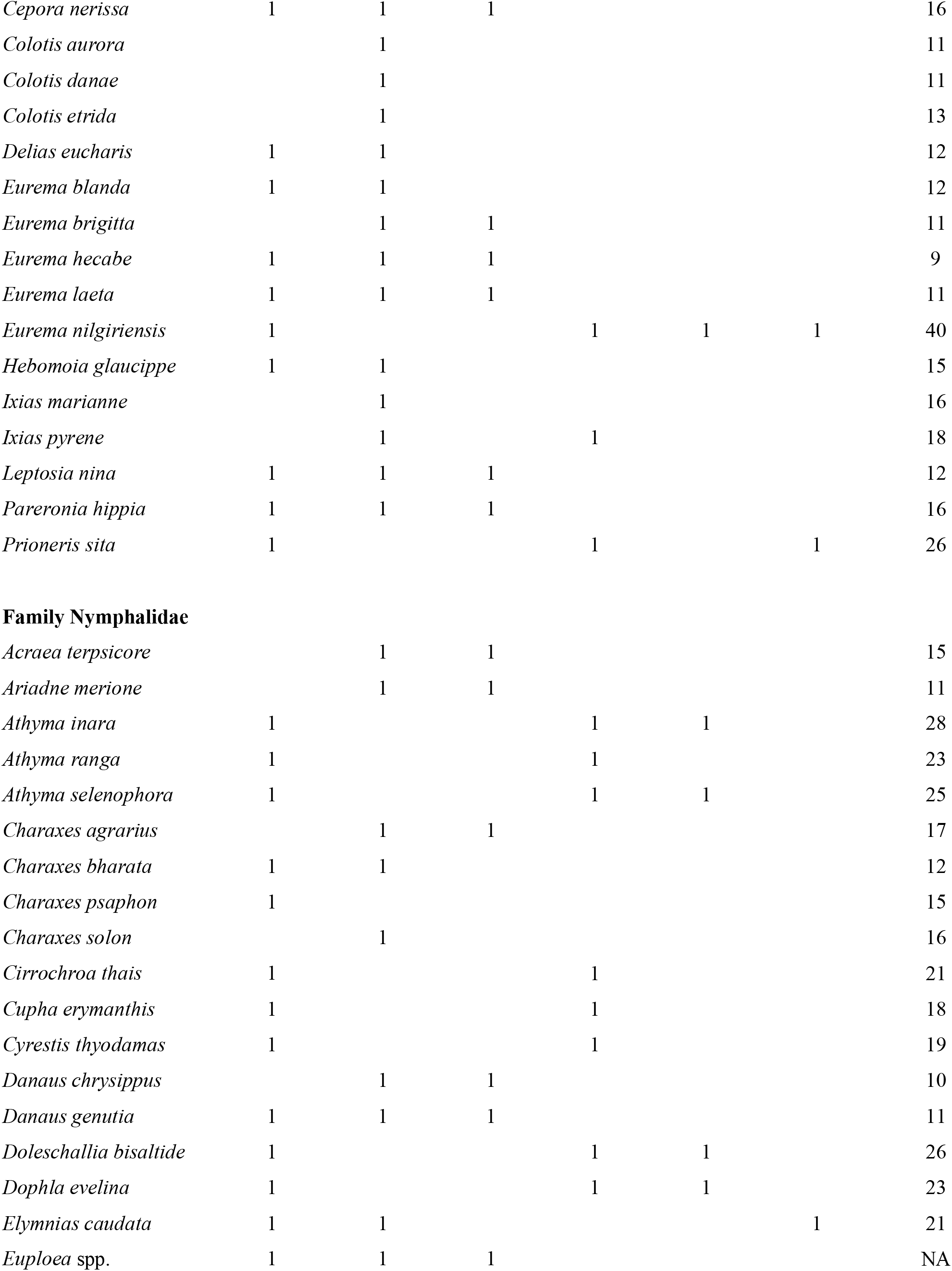
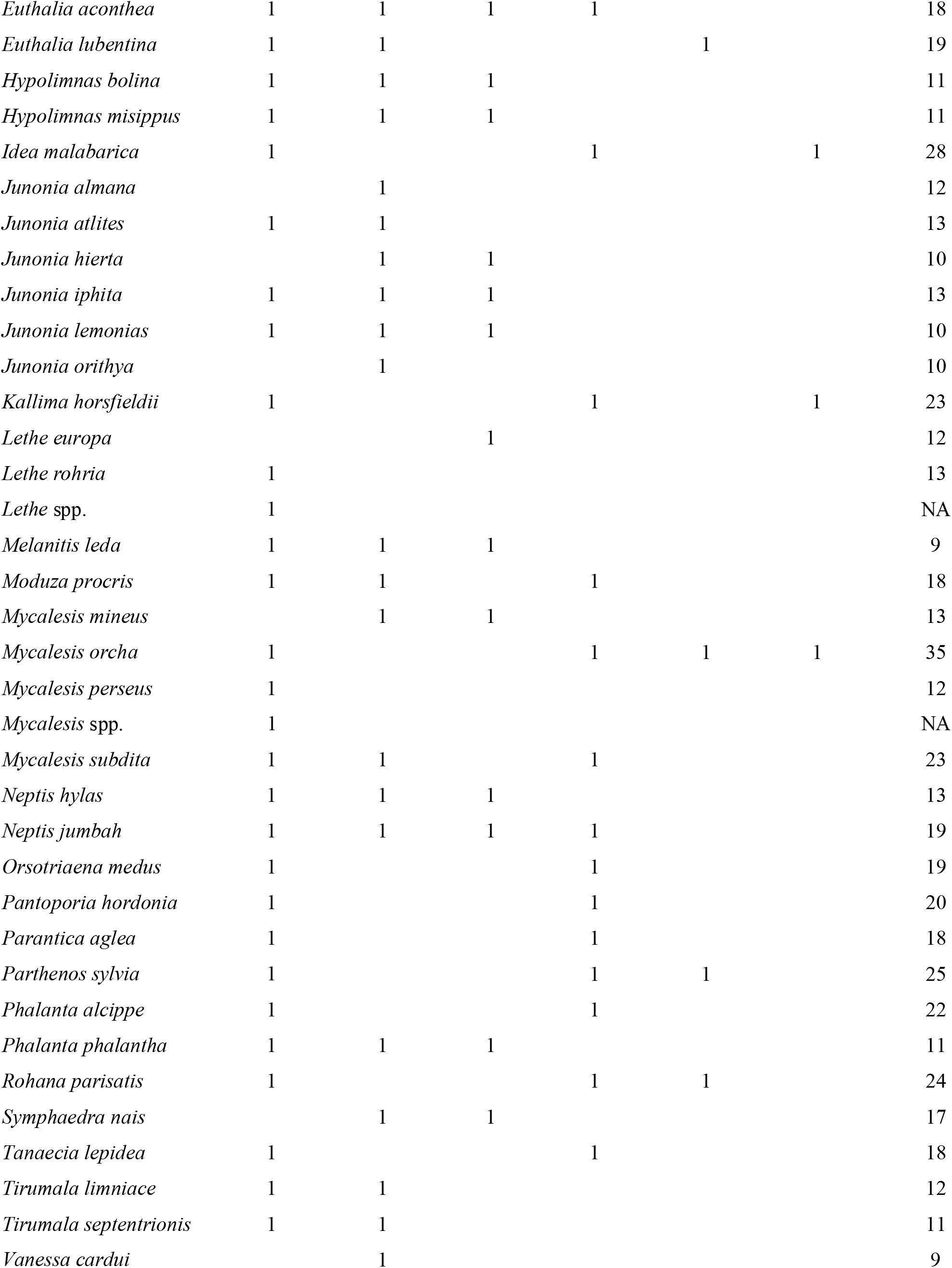
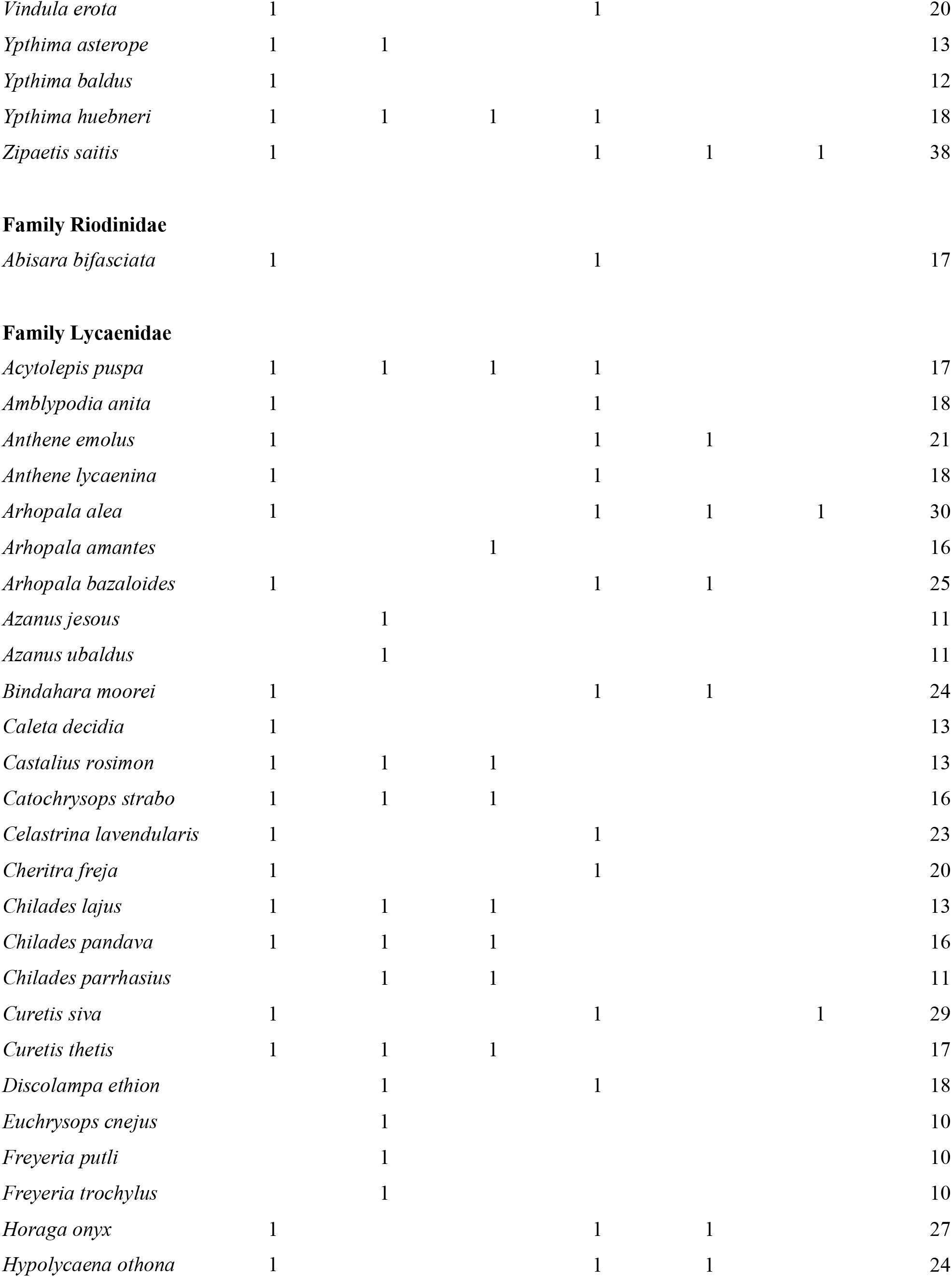
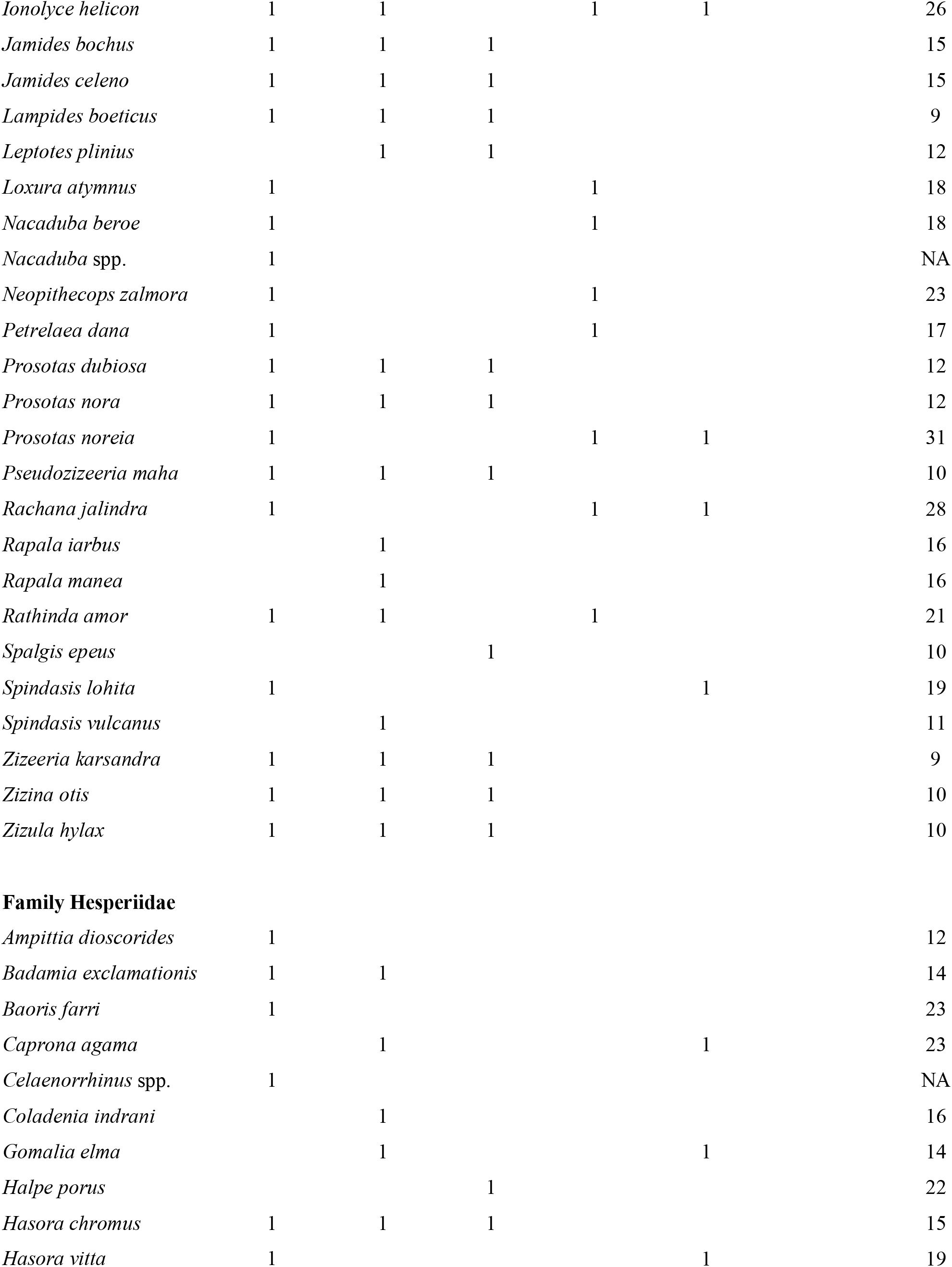
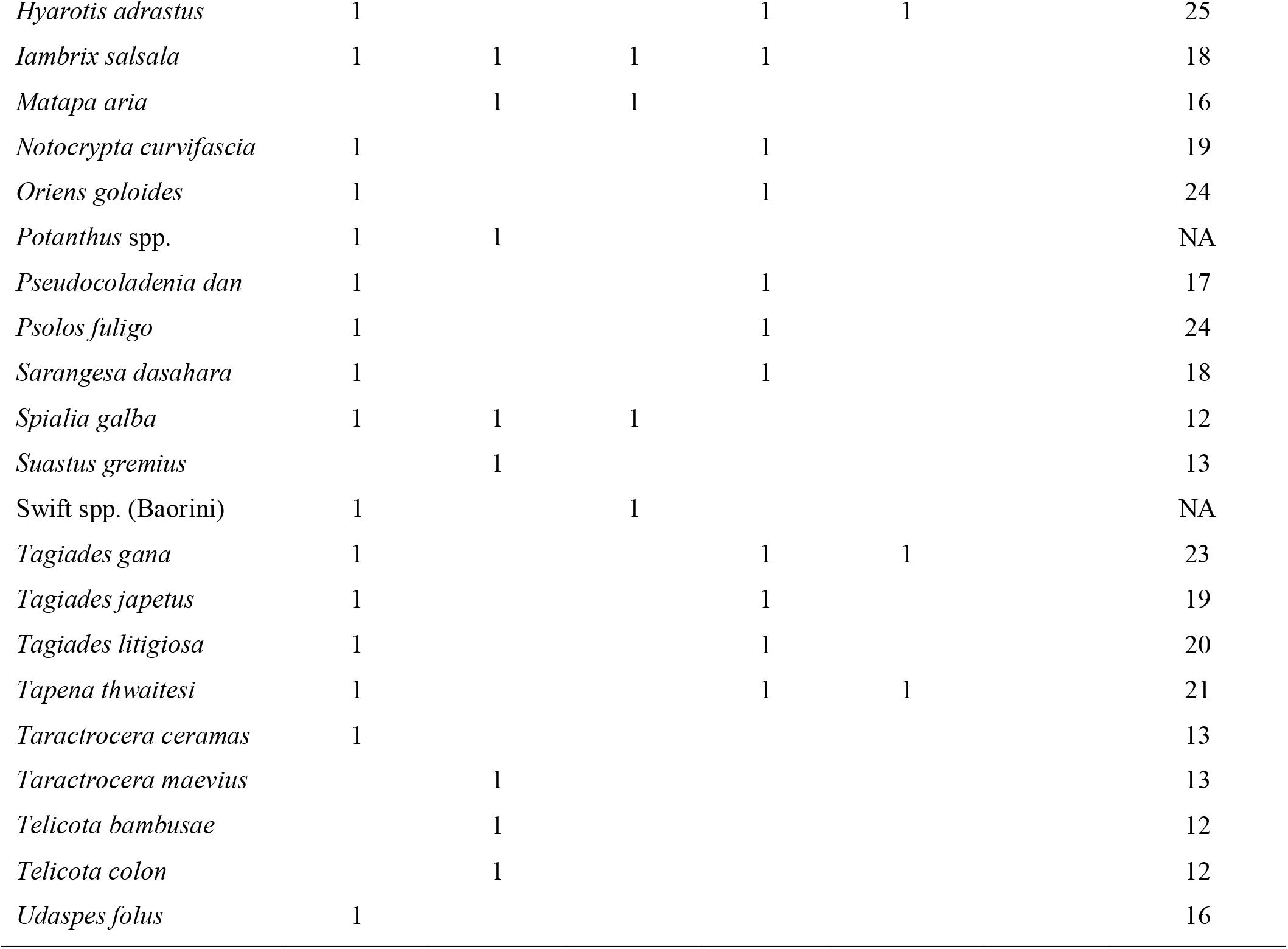
List of species observed in the study, indicating the habitat where they were observed and their habitat specialization, rarity and endemism. Species names follow Kunte et al. (2021), and conservation values are taken from Kunte (2008).

**Fig S1:**
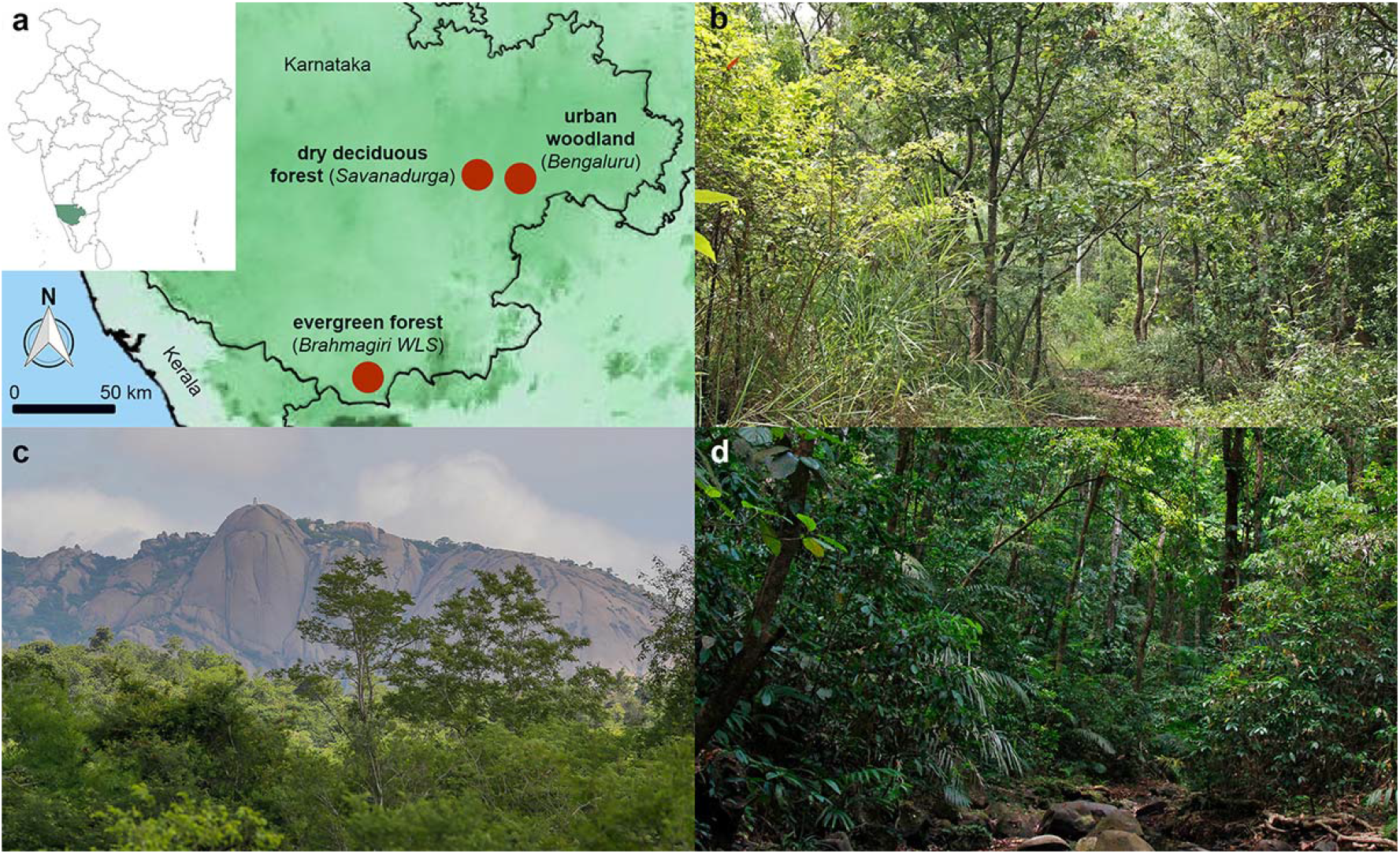
Butterfly habitats sampled in this study: (a) Locations of the three study sites. (b) Urban woodland in the Doresanipalya Forest Research Campus, Bengaluru, (c) Dry deciduous forest at Savanadurga, and (d) Evergreen forest in the Brahmagiri Wildlife Sanctuary, Kodagu.

## REFERENCES

Basset Y, Eastwood R, Sam L, et al (2013) Cross-continental comparisons of butterfly assemblages in tropical rainforests: implications for biological monitoring. Insect Conserv Divers 6:223–233. doi: 10.1111/j.1752-4598.2012.00205.x

Beneš J, Kepka P, Konvička M (2003) Limestone quarries as refuges for European xerophilous butterflies. Conserv Biol 17:1058–1069. doi: 10.1046/j.1523-1739.2003.02092.x

Bonebrake TC, Ponisio LC, Boggs CL, Ehrlich PR (2010) More than just indicators: A review of tropical butterfly ecology and conservation. Biol Conserv 143:1831–1841. doi: 10.1016/j.biocon.2010.04.044

Botham MS, Fernandez-Ploquin EC, Brereton T, et al (2015) Lepidoptera communities across an agricultural gradient: how important are habitat area and habitat diversity in supporting high diversity? J Insect Conserv 19:403–420. doi: 10.1007/s10841-015-9760-y

Brereton T, Roy DB, Middlebrook I, et al (2011) The development of butterfly indicators in the United Kingdom and assessments in 2010. J Insect Conserv 15:139–151. doi: 10.1007/s10841-010-9333-z

Brown KS (1997) Diversity, disturbance, and sustainable use of Neotropical forests: insects as indicators for conservation monitoring. J Insect Conserv 1:25–42. doi: 10.1023/A:1018422807610

Caldas A, Robbins RK (2003) Modified Pollard transects for assessing tropical butterfly abundance and diversity. Biol Conserv 110:211–219. doi: 10.1016/S0006-3207(02)00190-8

Chao A, Gotelli NJ, Hsieh TC, et al (2014) Rarefaction and extrapolation with Hill numbers: a framework for sampling and estimation in species diversity studies. Ecol Monogr 84:45–67. doi: 10.1890/13-0133.1

Chiarucci A, Bacaro G, Scheiner SM (2011) Old and new challenges in using species diversity for assessing biodiversity. Philos Trans R Soc B 366:2426–2437. doi: 10.1098/rstb.2011.0065

Clark TE, Samways MJ (1996) Dragonflies (Odonata) as indicators of biotope quality in the Kruger National Park, South Africa. J Appl Ecol 33:1001. doi: 10.2307/2404681

Devictor V, van Swaay C, Brereton T, et al (2012) Differences in the climatic debts of birds and butterflies at a continental scale. Nat Clim Chang 2:121–124. doi: 10.1038/nclimate1347

Fermon H, Waltert M, Mühlenberg M (2003) Movement and vertical stratification of fruit-feeding butterflies in a managed West African rainforest. J Insect Conserv 7:7–19. doi: 10.1023/A:1024755819790

Fourcade Y, WallisDeVries MF, Kuussaari M, et al (2021) Habitat amount and distribution modify community dynamics under climate change. Ecol Lett 24:950–957. doi: 10.1111/ele.13691

Glennie R, Buckland ST, Thomas L (2015) The effect of animal movement on line transect estimates of abundance. PLoS One 10:. doi: 10.1371/journal.pone.0121333

Gotelli NJ, Colwell RK (2001) Quantifying biodiversity: procedures and pitfalls in the measurement and comparison of species richness. Ecol Lett 4:379–391. doi: 10.1046/j.1461-0248.2001.00230.x

Henry EH, Haddad NM, Wilson J, et al (2015) Point-count methods to monitor butterfly populations when traditional methods fail: a case study with Miami blue butterfly. J Insect Conserv 19:519–529. doi: 10.1007/s10841-015-9773-6

Hill J, Hamer K, Tangah J, Dawood M (2001) Ecology of tropical butterflies in rainforest gaps. Oecologia 128:294–302. doi: 10.1007/s004420100651

Hsieh TC, Ma KH, Chao A (2016) iNEXT: an R package for rarefaction and extrapolation of species diversity (Hill numbers). Methods Ecol Evol 7:1451–1456. doi: 10.1111/2041-210X.12613

Isaac NJB, Cruickshanks KL, Weddle AM, et al (2011) Distance sampling and the challenge of monitoring butterfly populations. Methods Ecol Evol 2:585–594. doi: 10.1111/j.2041-210X.2011.00109.x

Jakubikova L, Kadlec T (2015) Butterfly bait traps versus zigzag walks: What is the better way to monitor common and threatened butterflies in non-tropical regions? J Insect Conserv 19:911–919. doi: 10.1007/s10841-015-9809-y

Kadlec T, Tropek R, Konvicka M (2012) Timed surveys and transect walks as comparable methods for monitoring butterflies in small plots. J Insect Conserv 16:275–280. doi: 10.1007/s10841-011-9414-7

Kéry M, Plattner M (2007) Species richness estimation and determinants of species detectability in butterfly monitoring programmes. Ecol Entomol 32:53–61. doi: 10.1111/j.1365-2311.2006.00841.x

Kim KC (1993) Biodiversity, conservation and inventory: why insects matter. Biodivers Conserv 2:191–214. doi: 10.1007/BF00056668

Kral-O’Brien KC, Antonsen AK, Hovick TJ, et al (2021) Getting the most from surveys: How method selection and method modification impact butterfly survey data. Ann Entomol Soc Am. doi: 10.1093/aesa/saab004

Kral K, Harmon J, Limb R, Hovick T (2018) Improving our science: the evolution of butterfly sampling and surveying methods over time. J Insect Conserv 22:1–14. doi: 10.1007/s10841-018-0046-z

Kremen C, Colwell RK, Erwin TL, et al (1993) Terrestrial arthropod assemblages: Their use in conservation planning. Conserv Biol 7:796–808. DOI: https://doi.org/10.1046/j.1523-1739.1993.740796.x

Kunte K (2008) The Wildlife (Protection) Act and conservation prioritization of butterflies of the Western Ghats, southwestern India. Curr Sci 94:729–735

Kunte K, Ravikanthachari N (2020) Butterflies of Bengaluru. 196

Kunte K, Sondhi S, Roy P (2020) Butterflies of India, v. 2.85

Leather SR, Basset Y, Hawkins BA (2008) Insect conservation: finding the way forward. Insect Conserv Divers 1:67–69. doi: 10.1111/j.1752-4598.2007.00005.x

Lewis OT, Basset Y (2007) Insect Conservation in Tropical Forests. In: Insect Conservation Biology. The Royal Entomological Society and CABI, Wallingford, pp 34–56

Lindenmayer DB, Likens GE (2010) The science and application of ecological monitoring. Biol Conserv 143:1317–1328. doi: 10.1016/j.biocon.2010.02.013

Mas AH, Dietsch T V (2003) An index of management intensity for coffee agroecosystems to evaluate butterfly species richness. Ecol Appl 13:1491–1501. DOI: https://doi.org/10.1890/01-5229

McDermott Long O, Warren R, Price J, et al (2017) Sensitivity of UK butterflies to local climatic extremes: which life stages are most at risk? J Anim Ecol 86:108–116. doi: 10.1111/1365-2656.12594

Mills SC, Oliver TH, Bradbury RB, et al (2017) European butterfly populations vary in sensitivity to weather across their geographical ranges. Glob Ecol Biogeogr 26:1374–1385. doi: 10.1111/geb.12659

Molleman F, Kop A, Brakefield PM, et al (2006) Vertical and temporal patterns of biodiversity of fruit-feeding butterflies in a tropical forest in Uganda. Biodivers Conserv 15:107–121. doi: 10.1007/s10531-004-3955-y

Nowicki P, Settele J, Henry P-Y, Woyciechowski M (2008) Butterfly monitoring methods: The ideal and the real world. Isr J Ecol Evol 54:69–88. doi: 10.1560/IJEE.54.1.69

Pardonnet S, Beck H, Milberg P, Bergman K-O (2013) Effect of tree-fall gaps on fruit-feeding Nymphalid butterfly assemblages in a Peruvian rain forest. Biotropica 45:612–619. DOI: https://doi.org/10.1111/btp.12053

Pellet J, Bried JT, Parietti D, et al (2012) Monitoring Butterfly Abundance: Beyond Pollard Walks. PLoS One 7:. doi: 10.1371/journal.pone.0041396

Pollard E, Yates TJ (1994) Monitoring butterflies for ecology and conservation: the British butterfly monitoring scheme, 1st edn. Springer Science & Business Media

Rohr JR, Mahan CG, Kim KC (2007) Developing a monitoring program for invertebrates: Guidelines and a case study. Conserv Biol 21:422–433. doi: 10.1111/j.1523-1739.2006.00578.x

Roy DB, Sparks TH (2000) Phenology of British butterflies and climate change. Glob Chang Biol 6:407–416. doi: 10.1046/j.1365-2486.2000.00322.x

Scheiner SM, Chiarucci A, Fox GA, et al (2011) The underpinnings of the relationship of species richness with space and time. Ecol Monogr 81:195–213. doi: 10.1890/10-1426.1

Schmeller DS, Henry P-Y, Julliard R, et al (2009) Advantages of volunteer-based biodiversity monitoring in Europe. Conserv Biol 23:307–316. doi: 10.1111/j.1523-1739.2008.01125.x

Sparrow HR, Sisk TD, Ehrlich PR, Murphy DD (1994) Techniques and guidelines for monitoring Neotropical butterflies. Conserv Biol 8:800–809. DOI: https://doi.org/10.1046/j.1523-1739.1994.08030800.x

Spitzer K, Novotny V, Tonner M, Leps J (1993) Habitat preferences, distribution and seasonality of the butterflies (Lepidoptera, Papilionoidea) in a montane tropical rain forest, Vietnam. J Biogeogr 20:109. doi: 10.2307/2845744

Taron D, Ries L (2015) Butterfly monitoring for conservation. In: Daniels JC (ed) Butterfly Conservation in North America. Springer, Netherlands, Dordrecht, pp 35–57

Thackeray SJ, Henrys PA, Hemming D, et al (2016) Phenological sensitivity to climate across taxa and trophic levels. Nature 535:241–245. doi: 10.1038/nature18608

Thomas J. (2005) Monitoring change in the abundance and distribution of insects using butterflies and other indicator groups. Philos Trans R Soc B Biol Sci 360:339–357. doi: 10.1098/rstb.2004.1585

Uehara-Prado M, Brown Jr KS, Freitas AVL (2007) Species richness, composition and abundance of fruit-feeding butterflies in the Brazilian Atlantic Forest: comparison between a fragmented and a continuous landscape. Glob Ecol Biogeogr 16:43–54. DOI: https://doi.org/10.1111/j.1466-8238.2006.00267.x

Van Dyck H, Van Strien AJ, Maes D, Van Swaay CAM (2009) Declines in Common, Widespread Butterflies in a Landscape under Intense Human Use. Conserv Biol 23:957–965. doi: 10.1111/j.1523-1739.2009.01175.x

van Strien AJ, van Swaay CAM, van Strien-van Liempt WTFH, et al (2019) Over a century of data reveal more than 80% decline in butterflies in the Netherlands. Biol Conserv 234:116–122. doi: 10.1016/j.biocon.2019.03.023

van Swaay CAM, Nowicki P, Settele J, van Strien AJ (2008) Butterfly monitoring in Europe: methods, applications and perspectives. Biodivers Conserv 17:3455–3469. doi: 10.1007/s10531-008-9491-4

Walpole MJ, Sheldon IR (1999) Sampling butterflies in tropical rainforest: an evaluation of a transect walk method. Biol Conserv 87:85–91. doi: 10.1016/S0006-3207(98)00037-8

Warren MS, Maes D, van Swaay CAM, et al (2021a) The decline of butterflies in Europe: Problems, significance, and possible solutions. Proc Natl Acad Sci 118:e2002551117. doi: 10.1073/pnas.2002551117

Warren MS, Maes D, van Swaay CAM, et al (2021b) The decline of butterflies in Europe: Problems, significance, and possible solutions. Proc Natl Acad Sci 118:. doi: 10.1073/pnas.2002551117

Zhang C, Harpke A, Kühn E, et al (2018) Applicability of butterfly transect counts to estimate species richness in different parts of the palaearctic region. Ecol Indic 95:735–740. doi: 10.1016/j.ecolind.2018.08.027

